# Postpartum cytokine shifts and IL-10–mediated immune suppression in malaria-infected primigravid women

**DOI:** 10.1101/2025.05.21.655391

**Authors:** Ousmane Traoré, Toussaint Rouamba, Serge Henri Zango, Hermann Sorgho, Innocent Valéa, Maminata Traoré-Coulibaly, Henk D. F. H. Schallig, Halidou Tinto

## Abstract

**Background:** According to the World Health Organization’s recent report, malaria remains a major health challenge during pregnancy and for postpartum women in endemic regions. While immune alterations during pregnancy are well characterized, postpartum cytokine dynamics and their impact on malaria susceptibility remain poorly defined. This study uniquely investigates how cytokine balance shifts, contribute to malaria susceptibility in primigravid women during the postpartum period.

**Methods:** A total of 33 Burkinabe women were enrolled at delivery and followed up at 1 and 3 months postpartum. Serum cytokine concentrations (IL-4, IL-6, IL-10, TNF-α, IFN-γ) were quantified by ELISA. Malaria infection was detected by PCR and microscopy. Statistical analyses included effect size calculations and cluster analyses to assess immune profiles.

**Results:** At delivery, 48.5% of women tested positive for malaria by PCR. Malaria-infected women had significantly elevated IL-10 levels and a decreased IL-6:IL-10 ratio compared with non-infected women (p = 0.005). This anti-inflammatory shift persisted into the during early postpartum period. Strong correlations were observed between IL-10 levels and malaria infection (σ = 0.9, p < 0.001). Of note, IL-4 also showed a significant effect, highlighting a complex immunoregulatory environment.

**Conclusion:** Our findings reveal, for the first time in a Sub-Saharan primigravid cohort, that an IL-10-dominant cytokine profile at delivery is strongly associated with postpartum malaria susceptibility. Modulating cytokine responses could represent a novel therapeutic approach to *improving* maternal health in malaria-endemic regions.

## INTRODUCTION

Malaria remains a major public health challenge, particularly among pregnant and postpartum women in endemic regions. Despite major advances in malaria control, maternal morbidity and mortality associated with *Plasmodium falciparum* infection persist, worsened by immune alterations during pregnancy and postpartum periods. Pregnancy induces a shift towards a Th2-dominant immune profile to protect the semi-allogeneic fetus (1–3). However, this immunological modulation makes pregnant women more vulnerable to infections, particularly malaria. After delivery, the postpartum period is characterized by progressive gradual immune reconstitution and rebalancing of Th1 and Th2 cytokine responses (4, 5). Recent studies suggest that the postpartum immune environment is distinct from that of pregnancy and non-pregnancy, but remains insufficiently understood, particularly with regard to malaria susceptibility Click or tap here to enter text.(6–8). Emerging data highlight the complex interplay between pro-inflammatory cytokines (e.g., IL-6, TNF-α, IFN-γ) and anti-inflammatory cytokines (e.g., IL-10, IL-4) in determining malaria outcomes (9, 10). Higher levels of IL-10 have been were reported to be involved in both protection against excessive inflammation and facilitation of parasite persistence (11, 12). However, few studies have longitudinally assessed cytokine dynamics specifically during the postpartum window, particularly in primigravid women who are at highest risk of malaria and its complications (13, 14). Furthermore, while studies have explored placental malaria and perinatal transmission, there is a paucity of research on how postpartum cytokine balances might influence new malaria infections or recrudescence (15, 16). The need for finer characterization of immunoregulatory mechanisms during postpartum recovery in malaria-endemic settings is pinpointed by recent immunological investigations (17, 18).. In the present study, we aim to address this critical gap by investigating the association between cytokine profiles and malaria susceptibility in primigravid women in during the early postpartum period in a high-transmission setting. We hypothesize that alterations ed in cytokine balance may predispose women to increased vulnerability to malaria after childbirth.

## METHODS

### Study Site

This study was conducted in the Nanoro Health and Demographic Surveillance System (HDSS) coverage area, located approximately 85 km northwest of Ouagadougou, the capital of Burkina Faso. Malaria transmission in this region is highly seasonal, with intense transmission during the rainy season, from June to October and a peak towards the end of the rainy season.

### Study design and sample collection

This study aimed to identify cytokine types associated with malaria infection during the postpartum period. It was nested within a larger clinical trial, the COSMIC study, which investigated the efficacy of different preventive strategies against malaria in pregnancy (trial registration numbers: ISRCTN372259296 and NCT01941564) (19). Participants in the current study were recruited from the control group of the COSMIC study, which followed the national policy of administering intermittent preventive treatment in pregnancy with sulfadoxine-pyrimethamine (IPTp-SP). Written informed consent was obtained from all participants, and they were followed up for 3 months after delivery. The characteristics of study participants are described elsewhere (20). Data on pregnancy background, including the number of IPTp-SP doses and history of malaria episodes, were obtained from the main study.

Ten milliliters of intravenous peripheral blood was collected in heparin tubes from each participant at delivery, and 1 and 3 months post-delivery. Blood samples were processed within 4 hours of collection, and sera were stored at −80°C until analysis. To prepare dried blood spots (DBS) for polymerase chain reaction (PCR) analysis, two 50-μl drops of blood were placed on filter paper (Whatman®) and allowed to dry completely for at least 4 hours at room temperature (25°C). To prepare plasma samples used for cytokine analysis, we carefully placed the heparinized tubes in a swing bucket centrifuge and spin them at 1800 rpm using a Hettich ROTANTA 460R centrifuge for 10 min at room temperature.

At each visit, women were systematically screened for malaria infection using the PfHRP2 rapid diagnostic test (RDT; SD-Bioline) following the manufacturer’s instructions for case management. The diagnostic test result was confirmed by light microscopy (LM) and PCR. Maternal peripheral blood was also collected at delivery to measure hemoglobin (Hb) level using a Hemocue (Hb 301, Sweden); anemia was defined as Hb ≤ 11g/dl.

### *Plasmodium falciparum* infectiondiagnosis by microscopy

Blood slides were prepared from samples collected at delivery, and at one and three months post-delivery. They were stained with Giemsa 3% for 45–60 min and examined by two independent microscopists blinded to RDT results. Parasite density was calculated against 200 leukocytes, or 500 leukocytes if less than 10 parasites were counted per 200 leukocytes. Slides were considered negative if no parasite was seen after examining 100 high-power fields. In case of discrepancies, a third independent reader’s opinion was required, and the final result was based on the two closest readings.

### Malaria diagnosis by nested-PCR

DNA was extracted from DBS using the QI Aamp DNA Extraction Mini-kit (QIAGEN, Valencia, CA, USA) and stored at −20°C until PCR amplification. A nested PCR for amplification of *Plasmodium falciparum* msp2 was performed in 25 µl reaction volumes, using 5 µl of DNA extract for the first round and 1 µl of first-round product for the second round with family-specific primers (21, 22). PCR products were visualized by ethidium bromide-stained agarose gel electrophoresis and UV transillumination, with fragment sizes estimated using Photo CaptMW (v11.01). End-point PCR cycling conditions included primary denaturation at 94°C for 5 minutes, followed by 36 cycles of denaturation at 94°C for 1 min, annealing at 58°C for 2 min, and extension at 72°C for 2 minutes, with a final extension at 72°C for 10 minutes.

### Cytokine ELISA

Commercially available cytokine ELISA kits for human samples were used to determine the levels of IL-4 (eBioscience BMS225/2), IL-10 (Invitrogen KHC0101), TNF-α (Invitrogen KHC3011), IL-6 (BioSource Europe KAC1261), and INF-γ (Invitrogen KAC1231). Each plate included a recombinant human cytokine standard curve and known positive and negative controls. All specimens were measured in duplicate, and the mean of the two values was taken. The lower limit of detection for each cytokine assay was 15 pg/mL.

### Placental malaria

Placental biopsy samples (2 cm × 2 cm × 1 cm) were collected from the maternal side at delivery, fixed in 10% neutral buffered formalin, and embedded in paraffin wax for histological analysis. Slides were stained with hematoxylin-eosin and read by trained microscopists. Placental infection was classified as acute (parasites present, malaria pigment absent), chronic (parasites and pigment present), past infection (no parasites, no pigment present), or no infection (no parasites or pigment) (23).

### Assessment of dystocia during labor

Dystocia during labor was diagnosed using various criteria in a partograph. A prolonged second stage of labor was documented, lasting more than 2 hours for nulliparous women (24), 3 hours with an epidural (25), and 1 hour for multiparous women. The reasons for prolonged labor were classified as mechanical or dynamic and could include weak contractions, fetal malposition, fetal macrosomia (24) cephalopelvic disproportion, and other medical or non-medical conditions. Other factors including maternal exhaustion, and pre-existing medical conditions were investigated. Blood loss during and immediately after delivery was estimated and documented, with categories of normal (< 500 mL for vaginal deliveries and < 1000 mL for cesarean sections) and excessive (indicating hemorrhage).

### Sample size and statistical analysis

The study included 33 primigravid women enrolled at delivery and followed up one and three months after delivery. The sample size (33 primigravida women) was determined based on the feasibility of the COSMIC study and the logistical constraints of participant follow-up. Although this sample size may be considered modest, it is consistent aligns with previous immunological studies examining cytokine responses in malaria-infected pregnant women (26). Given the observed differences in IL-10 levels between infected and non-infected women (σ = 0.9, p < 0.001), post-hoc power analysis indicates that our study had >80% power to detect significant differences in cytokine expression at an alpha level of 0.05. Future studies with larger cohorts will be needed to confirm these results in broader populations. Data were analyzed using Stata Statistical Software: version 15 (College Station, Texas). The Shapiro-Wilks W test was used to assess the normality of continuous variables, with logarithmic transformations applied where necessary. Spearman’s rho test was used to determine associations between continuous variables. Baseline cytokine levels in infected and uninfected women were compared using the Mann-Whitney test, unpaired Student’s t-test, or Kruskal-Wallis test.

To complement p-values, effect sizes were calculated using Cohen’s d to assess the magnitude of differences between cytokine levels in malaria-infected and non-infected women (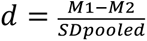; where 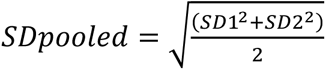). Effect sizes were classified as small (d = 0.2), medium (d = 0.5), and large (d ≥ 0.8). This analysis provided additional insights into the clinical relevance of cytokine differences beyond statistical significance. Serum cytokine concentrations were measured by enzyme-linked immunosorbent assay (ELISA) and analyzed using the “Cluster analysis” method (27), which requires a minimum of 5 to 10 individuals per group. Malaria infection status was determined by polymerase chain reaction (PCR) and microscopy. This dataset allowed a comprehensive examination of the relationships between cytokine profiles and malaria susceptibility.

## RESULTS

### Characteristics of the study population

A total of 33 primigravid women were enrolled in this study, with a median age of 19 years (range, 18 - 20.5years). At delivery, *P. falciparum* malaria infection was confirmed by PCR in 48.5% (n = 16) of women, while peripheral infection was determined by microscopy in 9.1% (n = 3). The mean parasite density at delivery was 540.9 [95% CI: 1–5,584] parasites/μL. Anemia was prevalent, with 60.6% (n = 20) of women anemic at delivery, but this rate decreased to 15.2% (n = 5) at one month postpartum before rising to 24.2% (n = 8) at three months postpartum. Placental malaria was detected in 23.8% (n = 5) of women at delivery (Table 1).

**Table 1.**
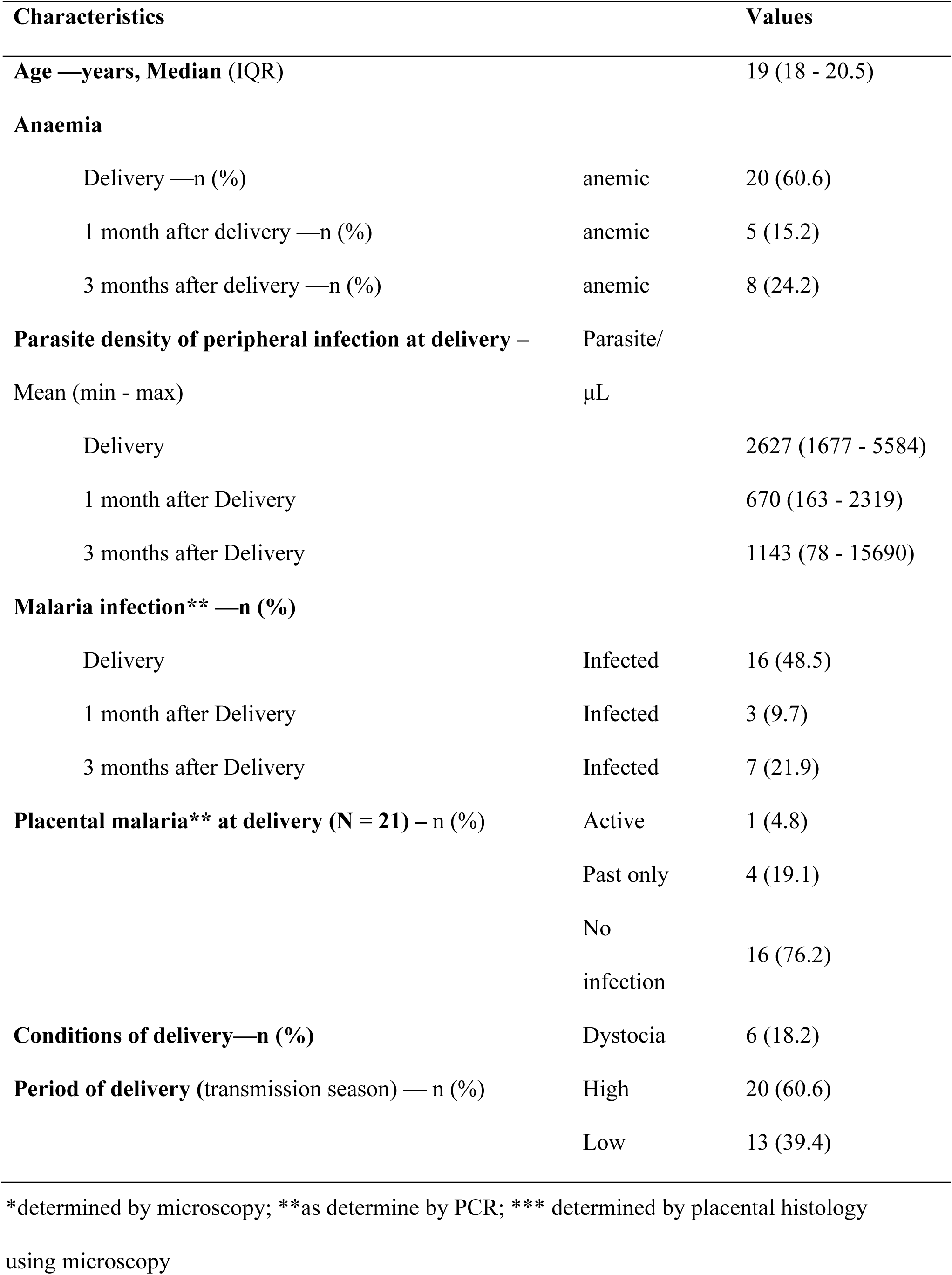
Characteristics of the study population.

### Cytokine distributions and correlations at delivery

At delivery, the levels of pro-inflammatory cytokines (IL-6, TNF-α) and anti-inflammatory cytokines (IL-4, IL-10) were analyzed to determine their correlations. Spearman’s rho test showed significant positive correlations between most cytokines. The strongest correlation was observed between IL-6 and IL-10 (σ = 0.8, p < 0.001) (Figure 1), followed by IL-10 and IL-4 (σ = 0.7, p < 0.001), and IL-6 and IL-4 (σ = 0.6, p < 0.001). Moderate correlations were found between IL-6 and TNF-α (σ = 0.5, p = 0.003), TNF-α and IL-4 (σ = 0.5, p = 0.002), and between IFN-γ and IL-4 (σ = 0.5, p = 0.007). In addition, IL-10 showed a moderate correlation with TNF-α (σ = 0.4, p = 0.013) and IFN-γ (σ = 0.4, p = 0.0169), and a weak correlation was observed between IL-6 and IFN-γ (σ = 0.3, p = 0.075).

**Figure 1.**
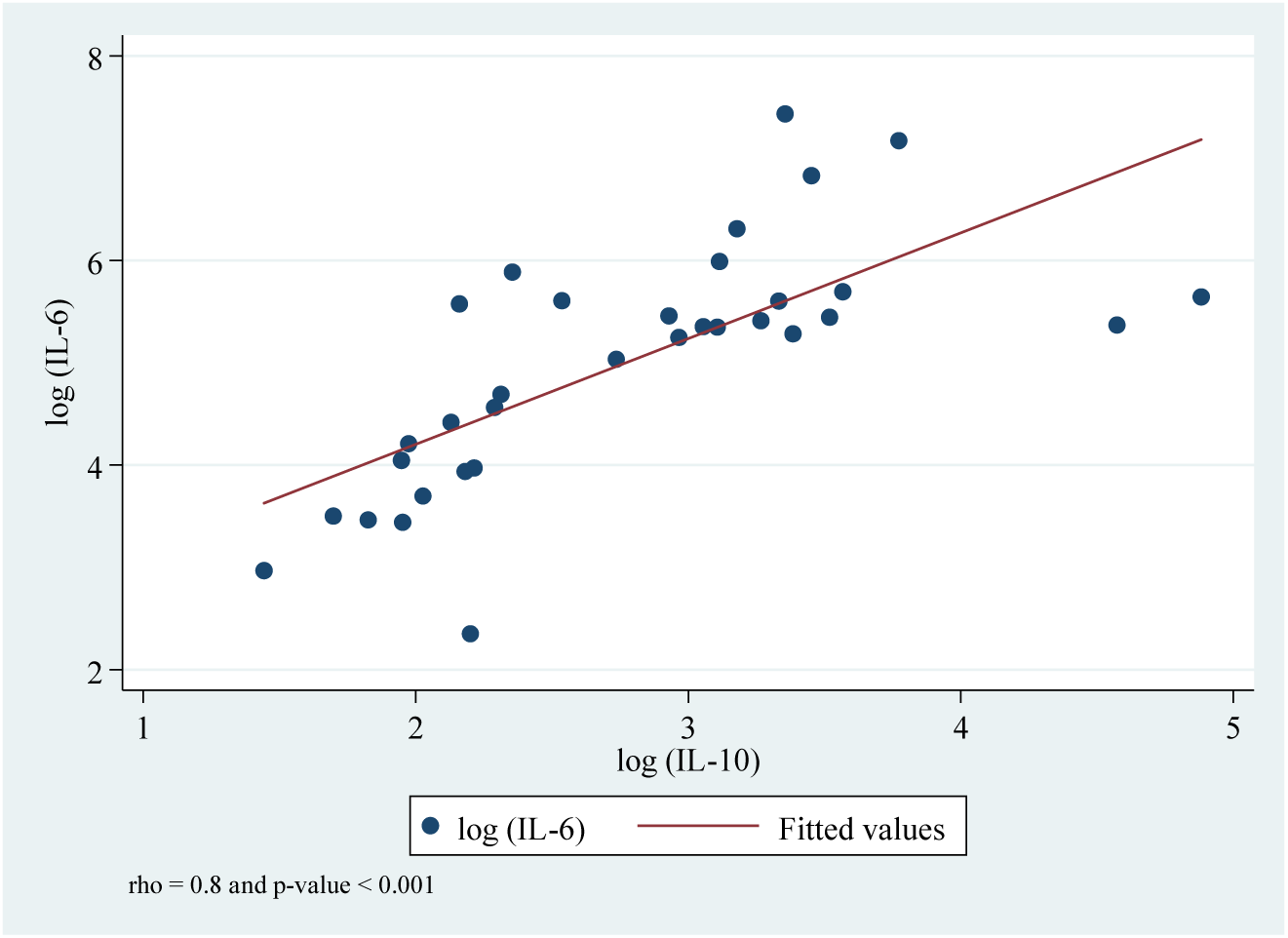
Relationship between serum IL-6 and IL-10 concentrations at delivery. The strongest correlation result following the Spearman’s correlation test coefficients is presented in this figure. Spearman’s correlation graded interpretation is defined as follows: (i) rho between 0.1 - 0.3 = weak; (ii) rho between 0.4 - 0.7 = moderate and (iii) rho between 0.8 - 1.0 = strong correlation.

To complement p-values, Cohen’s d was calculated to assess the effect sizes of cytokine differences between malaria-infected and non-infected women. IL-4 showed a medium-to-large effect size (d = 0.78), suggesting a notable difference between groups. IL-10 and IL-6 had medium effect sizes (d = 0.65 and d = 0.52, respectively), supporting their role in immune modulation during infection. In contrast, TNF-α (d = 0.08) and IFN-γ (d = 0.06) had small effects, indicating limited differences between infected and non-infected women (Table 2).

**Table 2.**
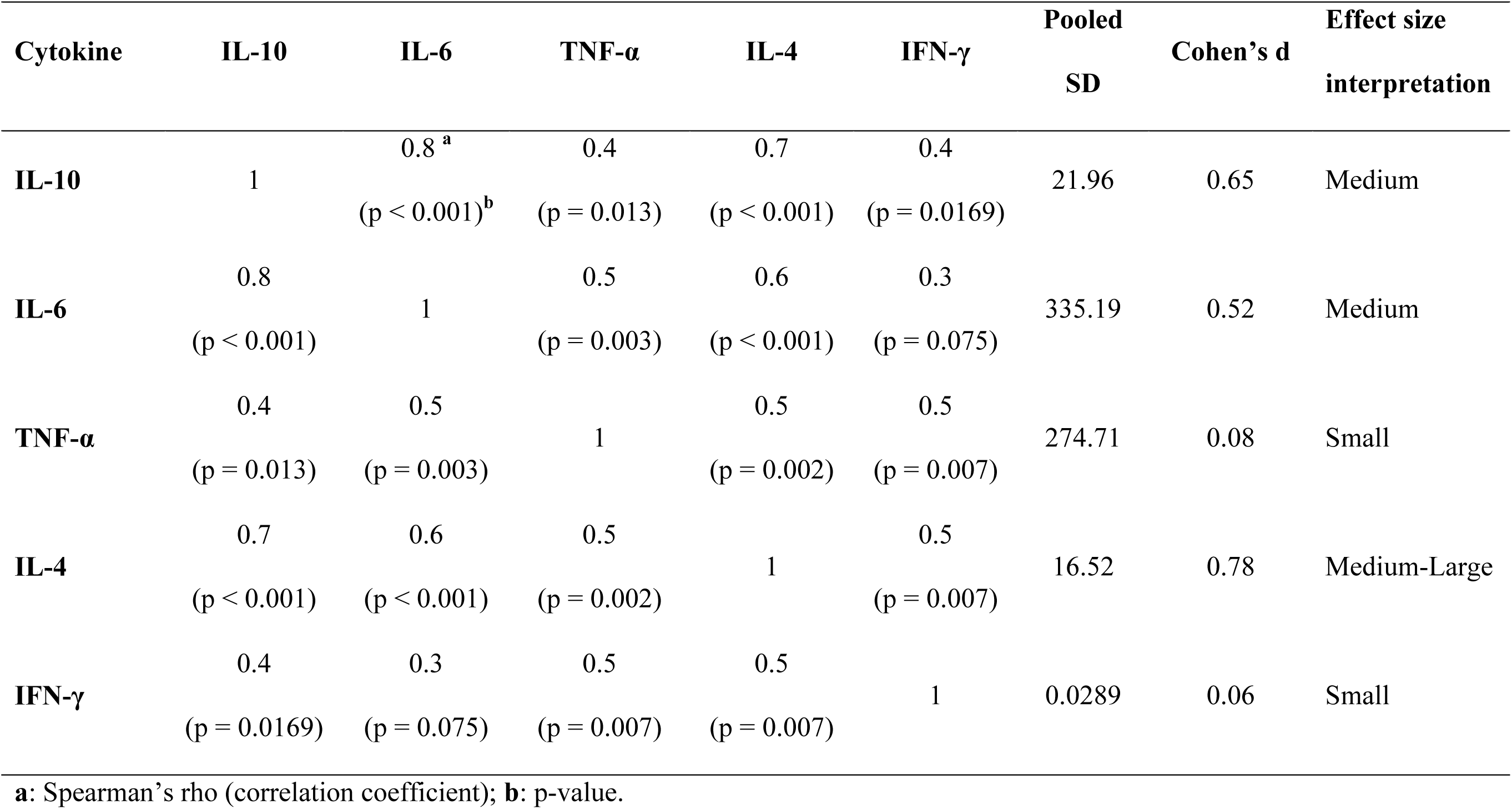
Correlations between cytokines concentrations (Spearman’s rho correlation coefficients), pooled SD, and effect sizes.

### Comparison of cytokine levels between infected and non-infected women at delivery

At delivery, serum levels of pro-inflammatory cytokines (IL-6: p < 0.001; TNF-α: p = 0.005; IFN-γ: p = 0.003) and anti-inflammatory cytokines (IL-4: p < 0.001; IL-10: p < 0.001) were significantly higher in malaria-infected women compared to non-infected women within the same cohort. Conversely, the IL-6: IL-10 ratio was significantly higher in non-infected women (p = 0.005) (Figures 2a-e).

**Figure 2:**
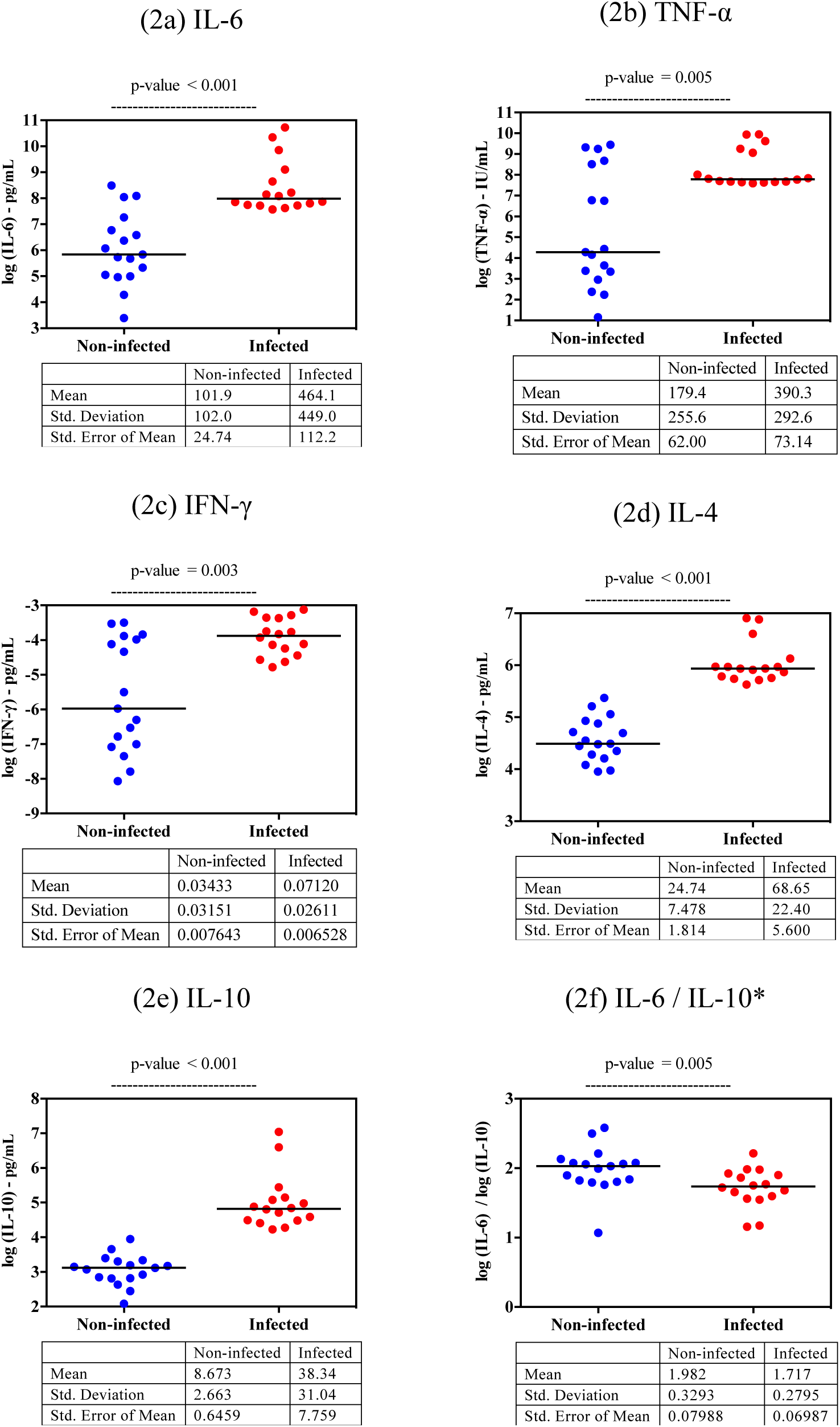
Comparison of cytokine levels between infected women and non-infected women at delivery. This figure illustrates the differences in cytokine levels between infected and non-infected women at delivery, highlighting key cytokines involved in the immune response to malaria. (2a) IL-6: Infected women show significantly higher IL-6 levels compared to non-infected women (p-value < 0.001), indicating a heightened pro-inflammatory response. (2b) TNF-α: TNF-α levels are also elevated in infected women (p-value = 0.005), further reflecting an increased inflammatory response. (2c) IFN-γ: There is a significant increase in IFN-γ levels in infected women (p-value = 0.003), suggesting enhanced activation of cellular immunity. (2d) IL-4: Infected women exhibit higher IL-4 levels (p-value < 0.001), indicating a concurrent anti-inflammatory response. (2e) IL-10: The levels of IL-10, a key anti-inflammatory cytokine, are significantly elevated in infected women (p-value < 0.001), which may contribute to immune modulation during malaria infection. (2f) IL-6 / IL-10 Ratio: The IL-6/IL-10 ratio is significantly lower in infected women (p-value = 0.005), indicating a relative dominance of anti-inflammatory over pro-inflammatory responses in these individuals.

### Pro- and anti-inflammatory cytokine balances during the postpartum period

The IL-6: IL-10 ratio was significantly higher in non-infected women at delivery [1.98 (1.81 - 2.15)] than in infected women [1.72 (1.57 - 1.87)], p = 0.019. However, this ratio did not differ significantly between infected and non-infected women, as was the PCR diagnosis at one and three months postpartum (Table 3).

**Table 3.** Pro- and anti-inflammatory cytokines balances at delivery, 1 month and 3 months after delivery.

|  | IL-6 to IL-10 ratios |  |  |
| --- | --- | --- | --- |
|  | Non-infected | Infected | p |
|  | n = 17 | n = 16 |  |
| **Delivery** | 1.98 (1.81 - 2.15) | 1.72 (1.57 - 1.87) | **0.019** |
|  | n = 28 | n = 3 |  |
| **1 Month after delivery** | 1.81 (1.54 - 2.09) | 1.56 (0.72 - 2.41) | 0.558 |
|  | n = 25 | n = 7 |  |
| **3 Months after delivery** | 1.88 (1.65 - 2.11) | 1.49 (1.01 - 1.97) | 0.105 |

p-values were calculated using t-test on log-transformed variables

### Comparison of cytokine levels during postpartum period

One month after delivery, infected women had significantly higher IL-10 levels (p = 0.012). Overall, cytokine levels were generally higher in infected women, although not all differences were statistically significant. At three months postpartum, cytokine levels were slightly higher in non-infected women with the exception of IL-4, which remained higher in infected women, but without significant differences (Table 4).

**Table 4.** Cytokines profiles (geometric mean (95% CI)) at 1 month and 3 months after delivery.

| Cytokines | Primiparous at 1 month after delivery |  |  | Primiparous at 3 months after delivery |  |  |
| --- | --- | --- | --- | --- | --- | --- |
|  | Non-infected | Infected | p | Non-infected | Infected | p |
|  | (n=28) | (n=3) |  | (n=25) | (n=7) |  |
| **IL-6 (pg/mL)** | 66.57<br>(34.21 - 129.57) | 238.43<br>(113.27 - 501.89) | 0.150 | 260.73<br>(103.75 - 655.23) | 74.53<br>(12.35 - 449.87) | 0.327 |
| **TNF-α (pg/mL)** | 54.01<br>(25.39 - 114.90) | 198.75<br>(97.43 - 405.44) | 0.242 | 104.81<br>(44.93 - 244.49) | 41.33<br>(9.60 - 177.83) | 0.494 |
| **IFN-γ (IU/mL)** | 0.05<br>(0.03 - 0.09) | 0.09<br>(0.06 - 0.17) | 0.664 | 0.10<br>(0.06 - 0.17) | 0.06<br>(0.02 - 0.16) | 0.121 |
| **IL-4 (pg/mL)** | 48.99<br>(37.47 - 64.08) | 77.35<br>(61.45 - 97.38) | 0.170 | 35.57<br>(26.35 - 48.00) | 31.85<br>(18.34 - 55.33) | 0.820 |
| **IL-10 (pg/mL)** | 10.56<br>(8.44 - 13.22) | 36.39<br>(7.89 - 167.72) | **0.012** | 19.56<br>(13.18 - 29.02) | 19.53<br>(6.40 - 59.58) | 0.715 |

\* p-values were calculated using t-test on log transformed values of parameters

### Relationships between cytokine concentrations and baseline characteristics

Cytokine levels at delivery were strongly correlated with malaria infection status, with IL-10 (σ = 0.9, p < 0.001) and IL-4 (σ = 0.9, p < 0.001) showing the strongest correlations. Parasite count also moderately correlated with IL-10 (σ = 0.5, p = 0.008). A negative correlation was observed between malaria infection status and the IL-6: IL-10 ratio (σ = −0.5, p = 0.004), and between parasite count and the IL-6: IL-10 ratio (σ = −0.4, p = 0.012) (Table 5).

**Table 5.** Relationships between serum cytokine concentrations and delivery characteristics.

| Cytokines | Women status at delivery |  |  |  |  |  |  |  |
| --- | --- | --- | --- | --- | --- | --- | --- | --- |
|  | Malaria |  | Dystocia |  | Anaemia |  | Parasitaemia |  |
| | $\sigma$ | p | $\sigma$ | p | $\sigma$ | p | | |
| **IL-6** | 0.7 | **<0.001** | 0.2 | 0.213 | -0.2 | 0.293 | 0.3 | 0.148 |
| **IL-10** | 0.9 | **<0.001** | 0.1 | 0.493 | 0.0 | 0.971 | 0.5 | **0.008** |
| **IL-4** | 0.9 | **<0.001** | 0.1 | 0.409 | 0.0 | 0.857 | 0.2 | 0.288 |
| **IFN-γ** | 0.5 | **0.002** | 0.2 | 0.359 | -0.1 | 0.614 | 0.3 | 0.135 |
| **TNF-α** | 0.5 | **0.004** | 0.0 | 0.784 | -0.1 | 0.516 | 0.1 | 0.457 |
| **IL-6: IL-10** | -0.5 | **0.004** | 0.0 | 0.820 | 0.0 | 0.943 | -0.4 | **0.012** |

## DISCUSSION

This study showed that malaria-infected women had significantly higher levels of both pro-inflammatory (IL-6, TNF-α, IFN-γ) and anti-inflammatory (IL-4, IL-10) cytokines at delivery compared to non-infected women. The IL-6 : IL-10 ratio was notably lower in infected women, suggesting a dominance of anti-inflammatory responses during malaria infection. These cytokine balances shifted during the postpartum period, with non-infected women showing higher IL-6: IL-10 ratios at 1 and 3 months postpartum.

The elevated levels of IL-10 in malaria-infected women at delivery align with previous research highlighting the critical role of cytokines in modulating immune responses to *Plasmodium falciparum* infection. High IL-10 levels have been associated with reduced severity of malaria symptoms by moderating the inflammatory response (28).

However, our findings suggest that excessive IL-10 production may suppress effective pro-inflammatory responses necessary for parasite elimination, potentially contributing to persistent infection and adverse pregnancy outcomes (26, 28).

The lower IL-6 : IL-10 ratio observed in malaria-infected women at delivery indicates an anti-inflammatory bias, which might allow the parasite to evade immune clearance (28, 29). This aligns with studies demonstrating that IL-10 inhibits Th1-mediated immune responses, limiting the activation of macrophages and dendritic cells necessary for pathogen clearance (30). The flowchart in Figure 4, illustrates the proposed cytokine balance shift, depicting how IL-10 overproduction may facilitate *Plasmodium falciparum* persistence while mitigating severe inflammation.

**Figure 3:**
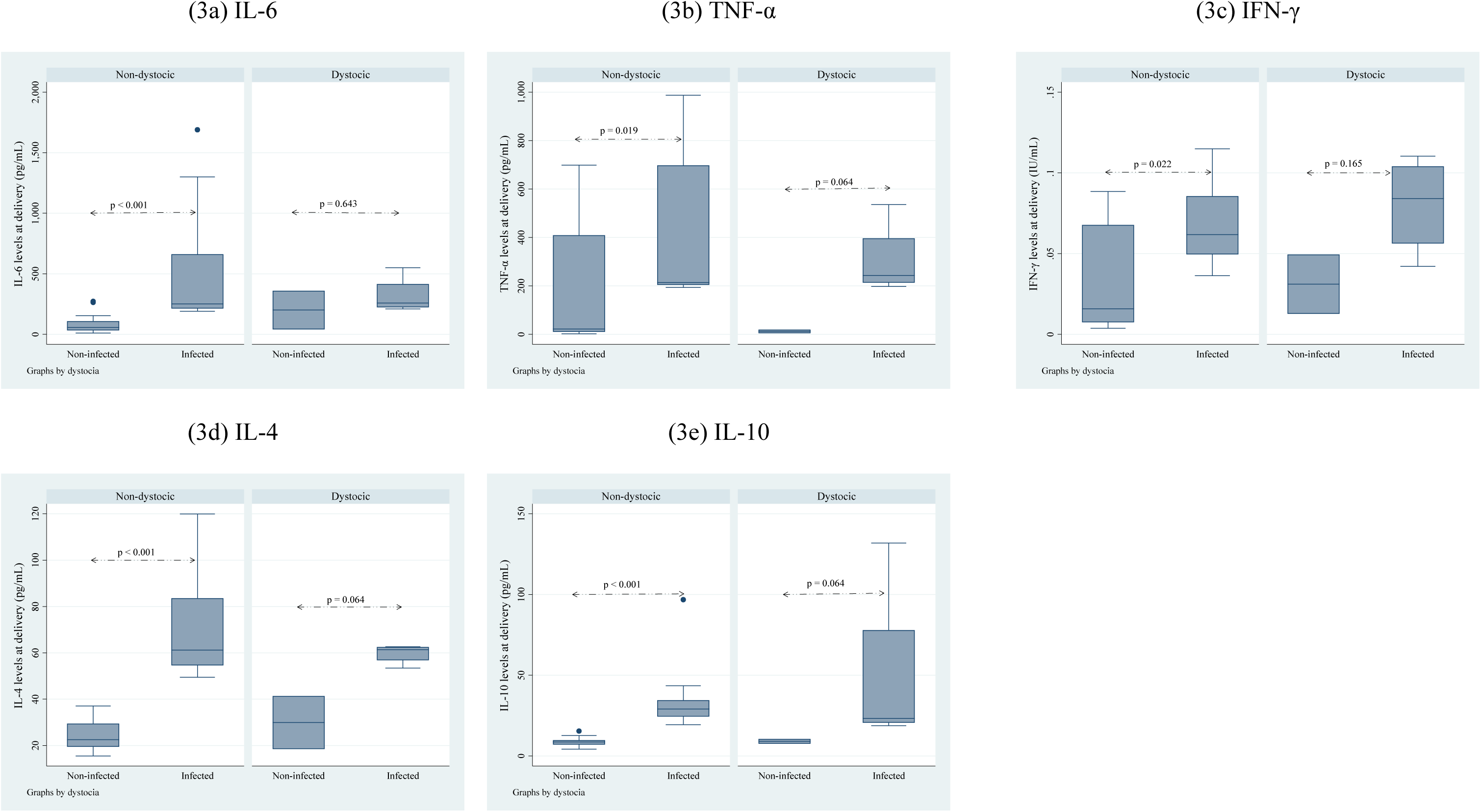
Assessment of the effect of dystocia on cytokine levels at delivery. This illustration reports the effect of dystocia on cytokine levels at delivery in women infected and non-infected by *P. falciparum*. IL-6 levels (3a) are significantly higher in infected women with dystocia compared to non-infected women (p < 0.001). TNF-α (3b) also shows a significant increase in the infected group (p < 0.05). IFN-γ levels (3c) are elevated in infected women, but the difference is not statistically significant. Both IL-4 (3d) and IL-10 (3e) levels are significantly higher in infected women, with IL-10 showing the most substantial difference (p < 0.001). This suggests that dystocia impacts cytokine levels, particularly in infected women, indicating an enhanced inflammatory response.

**Figure 4:**
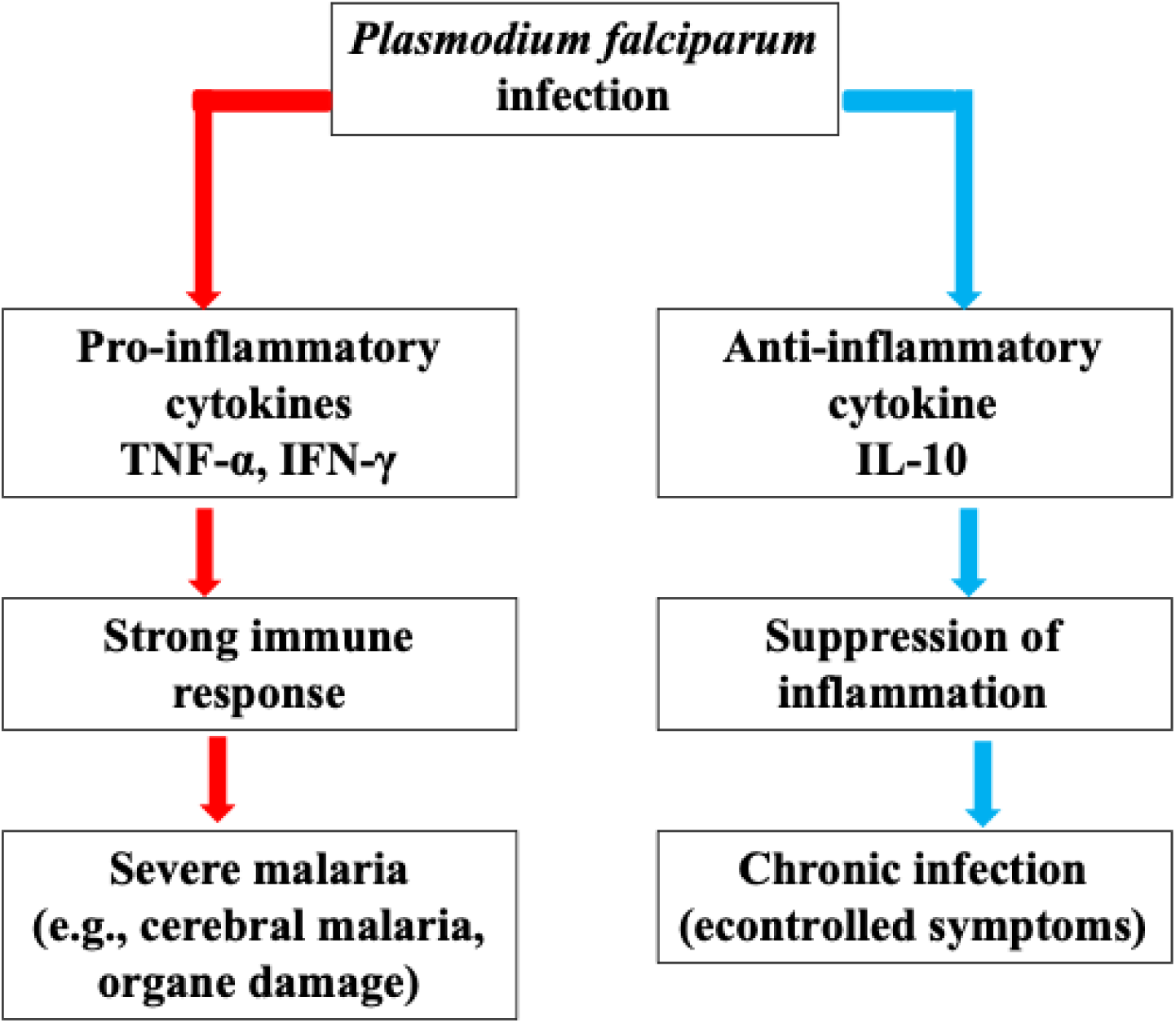
Cytokine Balance in *Plasmodium falciparum* Infection. The immune response to *Plasmodium falciparum* infection involves a delicate balance between pro-inflammatory and anti-inflammatory cytokines. Pro-inflammatory cytokines such as TNF-α and IFN-γ trigger a strong immune response, which is essential for parasite clearance but also increases the risk of severe inflammation and tissue damage. In contrast, IL-10 suppresses excessive inflammation, reducing immune-mediated pathology but facilitating parasite persistence. The clinical outcome depends on this cytokine balance: a dominant pro-inflammatory response can lead to severe malaria (e.g., cerebral malaria, organ damage), whereas a high IL-10 response may promote chronic infection with controlled symptoms.

The postpartum period represents a critical window for immune reconstitution as the maternal immune system transitions from pregnancy-induced tolerance to a more balanced state (10). The higher IL-6: IL-10 ratios in non-infected women during this period suggest a recovery of pro-inflammatory responses, essential for combating infections (31). This shift in cytokine balance towards a pro-inflammatory state might explain the reduced susceptibility to malaria observed in non-infected women as they recover from pregnancy. Furthermore, regulatory T cells (Tregs) are linked to immune modification during malaria infection.

Research indicates that Tregs producing IL-10 reduce antigen-specific immune responses, so promoting chronic parasitemia (32). The spread of these cells in postpartum malaria-infected women might help to explain extended susceptibility.

Physical factors such as dystocia were also considered in this study. Although dystocia was associated with higher cytokine levels in infected women, it did not significantly alter the overall cytokine balance between infected and non-infected groups. This suggests that while physical stressors contribute to inflammation, malaria infection itself is the primary driver of cytokine changes (33, 34).

The use of the cluster analysis approach ensured sufficient statistical power to detect significant differences in cytokine levels between infected and non-infected women (27) Our findings highlight the complex relationship between inflammatory cytokines and malaria infection status during postpartum, suggesting that cytokine profiles may serve as biomarkers for malaria pathogenesis in this population. Therapeutic approaches targeting cytokine modulation could offer novel strategies to improve clinical outcomes. IL-4, for instance, has been shown to reduce parasitaemia, cerebral malaria pathology, and mortality by promoting parasite clearance while reducing brain inflammation (35). Additionally, rosiglitazone, a PPARγ agonist, enhances phagocytic clearance of parasitized erythrocytes and reduces inflammatory responses in experimental cerebral malaria by inhibiting specific signaling pathways(36).

Furthermore, elevated IFN-γ and TNF-α levels have been associated lower fever and malaria risk, while IL-12 production plays a role in reducing parasitemia. Conversely, an imbalanced proinflammatory-to-anti-inflammatory cytokine ratio may increase the risk of severe malaria symptoms (37). These findings underscore the potential of targeting cytokine profiles as a therapeutic avenue.

Our study highlights key insights, but further research is needed to identify the cellular sources of IL-10 and its role in postpartum malaria immunity. Future research should include:

i. PBMC stimulation assays to verify whether IL-10 suppresses protective inflammatory responses.
ii. Treg profiling by flow cytometry to examine the role of regulatory T cells in malaria persistence.
iii. RNA-seq or proteomic analyses to study signaling pathways downstream of cytokine modulation.
iv. Clinical trials with IL-10 inhibitors to test their effectiveness in antimalarial immunity without increasing inflammation.

Furthermore, studying the impact of interventions aimed at modulating cytokine responses could offer promising avenues for improving maternal and fetal health in malaria-endemic regions.

## CONCLUSION

This study shows that women infected with malaria-postpartum exhibit a distinct cytokine profile, with elevated IL-10 and a suppressed IL-6:IL-10 ratio at delivery. These findings suggest an anti-inflammatory bias that may impair immune clearance and promote parasite persistence. In contrast, non-infected women regained their pro-inflammatory responses after delivery. The observed cytokine shifts highlight the potential of immune profiling as a biomarker for malaria risk and support further research into immunomodulatory therapies for postpartum malaria.

## Acknowledgements

We are grateful to the COSMIC trial participants and the field staff of the Clinical Research Unit of Nanoro for their contribution to the study completion. We would like to thank the nulliparous women recruited in the HDSS catchment area.

## Declarations

### Ethical approval

This study received ethical approval from the institutional review board of Centre Muraz/IRSS (Reference A007-2014/CEI-CM dated 12^th^ February 2014).

### Availability of Data and Materials

The data supporting the conclusions of this article are available upon request from the corresponding author.

### Authors’ contributions

OT conceived, designed and performed the experiments, OT and TR analyzed the data. SHZ, HS, IV, MTC supervised the fieldwork. OT drafted the manuscript. SHZ, HS, IV, MTC, HDFHS, and HT revised the manuscript. All authors contributed to improving the draft and read and approved the final manuscript.

## REFERENCES

1. Halonen M, Lohman IC, Stern DA, Spangenberg A, Anderson D, Mobley S, Ciano K, Peck M, Wright AL. 2009. Th1/Th2 Patterns and Balance in Cytokine Production in the Parents and Infants of a Large Birth Cohort. J Immunol 182:3285–3293.

2. Denney JM, Nelson EL, Wadhwa PD, Waters TP, Mathew L, Chung EK, Goldenberg RL, Culhane JF. 2011. Longitudinal modulation of immune system cytokine profile during pregnancy. Cytokine 53:170–177.

3. Maestre A, Carmona-Fonseca J. 2014. Immune responses during gestational malaria: a review of the current knowledge and future trend of research. J Infect Dev Ctries 8:391–402.

4. Mclean ARD, Ataide R, Simpson JA, Beeson JG, Fowkes FJI. 2015. Malaria and immunity during pregnancy and postpartum: a tale of two species. Parasitology 142:999– 1015.

5. D’Ambruoso L, Byass P, Qomariyah SN, Ouédraogo M. 2010. A lost cause? Extending verbal autopsy to investigate biomedical and socio-cultural causes of maternal death in Burkina Faso and Indonesia. Soc Sci Med 71:1728–1738.

6. Nguyen-Dinh P, Steketee Richard W, Greenberg Alan E, Wirima Jack J, Mulenda O, Williams Sharyon B. 1988. RAPID SPONTANEOUS POSTPARTUM CLEARANCE OF PLASMODIUM FALCIPARUM PARASITAEMIA IN AFRICAN WOMEN. Lancet 332:751–752.

7. Massougbodji A, Doritchamou J, Briand V, Agbowai C, Bottero J, Cot M. 2011. Spontaneous Postpartum Clearance of Plasmodium falciparum Parasitemia in Pregnant Women, Benin. Am J Trop Med Hyg 84:267–269.

8. Boel ME, Rijken MJ, Brabin BJ, Nosten F, McGready R. 2012. The epidemiology of postpartum malaria: a systematic review 11:114.

9. Diagne N, Rogier C, Sokhna CS, Tall A, Fontenille D, Roussilhon C, Spiegel A, Trape J-F. 2000. Increased Susceptibility to Malaria during the Early Postpartum Period. New Engl J Medicine 343:598–603.

10. Ramharter M, Grobusch MP, Kießling G, Adegnika AA, Möller U, Agnandji STM, Kramer M, Schwarz N, Kun JFJ, Oyakhirome S, Issifou S, Borrmann S, Lell B, Mordmüller B, Kremsner PG. 2005. Clinical and Parasitological Characteristics of Puerperal Malaria. J Infect Dis 191:1005–1009.

11. Deloron P, Lombard PR, Ringwald P, Wallon M, Niyongabo T, Aubry P, Dayer JM, Peyron F. 1994. Plasma levels of TNF-alpha soluble receptors correlate with outcome in human falciparum malaria. Eur cytokine Netw 5:331–6.

12. Langhorne J, Ndungu FM, Sponaas A-M, Marsh K. 2008. Immunity to malaria: more questions than answers. Nat Immunol 9:725–732.

13. Dingermann T. 2006. The Interferons: Characterization and Application. Von Anthony Meager (Hrsg.). Pharm unserer Zeit 35:464–464.

14. Lotze M, Hamilton J. 2003. The Cytokine Handbook 927–958.

15. Riley EM. 1996. The role of MHC- and non-MHC-associated genes in determining the human immune response to malaria antigens. Parasitology 112:S39–S51.

16. Schofield L, Grau GE. 2005. Immunological processes in malaria pathogenesis. Nat Rev Immunol 5:722–735.

17. Sacks GP, Studena K, Sargent IL, Redman CWG. 1998. Normal pregnancy and preeclampsia both produce inflammatory changes in peripheral blood leukocytes akin to those of sepsis. Am J Obstet Gynecol 179:80–86.

18. Healy LL, Cronin JG, Sheldon IM. 2014. Endometrial cells sense and react to tissue damage during infection of the bovine endometrium via interleukin 1. Sci Rep 4:7060.

19. Scott S, Mens PF, Tinto H, Nahum A, Ruizendaal E, Pagnoni F, Grietens KP, Kendall L, Bojang K, Schallig H, D’Alessandro U. 2014. Community-based scheduled screening and treatment of malaria in pregnancy for improved maternal and infant health in The Gambia, Burkina Faso and Benin: study protocol for a randomized controlled trial. Trials 15:340.

20. Ousmane T, Hermann S, Isidore Y W, Toussaint R, S S Guillaume, Innocent V, Maminata T-C, Susana S, F M Petra, Henk S. 2018. Naturally acquired antibody to DBL5 and ID1-ID2a dynamics in primigravid women during postpartum in a rural setting of Burkina Faso. African Journal of Immunology Research 5:453–462.

21. Wooden J, Kyes S, Sibley CH. 1993. PCR and strain identification in Plasmodium falciparum. Parasitol Today 9:303–305.

22. Singh RP, Nie X, Singh M, Coffin R, Duplessis P. 2002. Sodium sulphite inhibition of potato and cherry polyphenolics in nucleic acid extraction for virus detection by RT-PCR. J Virol Methods 99:123–131.

23. Bulmer JN, Rasheed FN, Francis N, Morrison L, Greenwood BM. 1993. Placental malaria. I. Pathological classification. Histopathology 22:211–218.

24. (College) AC of O and G, Medicine S for M-F, Caughey AB, Cahill AG, Guise J-M, Rouse DJ. 2014. Safe prevention of the primary cesarean delivery. Am J Obstet Gynecol 210:179–193.

25. Anim-Somuah M, Smyth RM, Cyna AM, Cuthbert A. 2018. Epidural versus non-epidural or no analgesia for pain management in labour. Cochrane Database Syst Rev 2018:CD000331.

26. Megnekou R, Staalsoe T, Hviid L. 2013. Cytokine response to pregnancy-associated recrudescence of Plasmodium berghei infection in mice with pre-existing immunity to malaria. Malar J 12:387.

27. Genser B, Cooper PJ, Yazdanbakhsh M, Barreto ML, Rodrigues LC. 2007. A guide to modern statistical analysis of immunological data. BMC Immunol 8:27.

28. Fried M, Muga RO, Misore AO, Duffy PE. 1998. Malaria elicits type 1 cytokines in the human placenta: IFN-gamma and TNF-alpha associated with pregnancy outcomes. J Immunol (Baltim, Md : 1950) 160:2523–30.

29. Chandrasiri UP, Randall LM, Saad AA, Bashir AM, Rogerson SJ, Adam I. 2014. Low Antibody Levels to Pregnancy-specific Malaria Antigens and Heightened Cytokine Responses Associated With Severe Malaria in Pregnancy. J Infect Dis 209:1408–1417.

30. Bouyou-Akotet MK, Issifou S, Meye JF, Maryvonne K, Ngou-Milama E, Luty AJF, Kremsner PG, Mavoungou E. 2004. Depressed Natural Killer Cell Cytotoxicity against Plasmodium falciparum–Infected Erythrocytes during First Pregnancies. Clin Infect Dis 38:342–347.

31. Achidi EA, Apinjoh TO, Titanji VPK. 2007. Malaria parasitemia and systemic cytokine bias in pregnancy. Int J Gynecol Obstet 97:15–20.

32. Kumar R, Ng S, Engwerda C. 2019. The Role of IL-10 in Malaria: A Double Edged Sword. Front Immunol 10:229.

33. Scaffidi P, Misteli T, Bianchi ME. 2002. Release of chromatin protein HMGB1 by necrotic cells triggers inflammation. Nature 418:191–195.

34. Tian J, Avalos AM, Mao S-Y, Chen B, Senthil K, Wu H, Parroche P, Drabic S, Golenbock D, Sirois C, Hua J, An LL, Audoly L, Rosa GL, Bierhaus A, Naworth P, Marshak-Rothstein A, Crow MK, Fitzgerald KA, Latz E, Kiener PA, Coyle AJ. 2007. Toll-like receptor 9–dependent activation by DNA-containing immune complexes is mediated by HMGB1 and RAGE. Nat Immunol 8:487–496.

35. Wu X, Thylur RP, Dayanand KK, Punnath K, Norbury CC, Gowda DC. 2021. IL-4 Treatment Mitigates Experimental Cerebral Malaria by Reducing Parasitemia, Dampening Inflammation, and Lessening the Cytotoxicity of T Cells. J Immunol 206:118–131.

36. Serghides L, Patel SN, Ayi K, Lu Z, Gowda DC, Liles WC, Kain KC. 2009. Rosiglitazone Modulates the Innate Immune Response to Plasmodium falciparum Infection and Improves Outcome in Experimental Cerebral Malaria. J Infect Dis 199:1536–1545.

37. Dodoo D, Omer FM, Todd J, Akanmori BD, Koram KA, Riley EM. 2002. Absolute Levels and Ratios of Proinflammatory and Anti-inflammatory Cytokine Production In Vitro Predict Clinical Immunity to Plasmodium falciparum Malaria. J Infect Dis 185:971–979.

